# Subcellular plant carbohydrate metabolism under elevated temperature

**DOI:** 10.1101/2024.10.31.621406

**Authors:** Charlotte Seydel, Martin Heß, Laura Schröder, Andreas Klingl, Thomas Nägele

**Affiliations:** LMU München, Faculty of Biology, Plant Development, Großhaderner Str. 2-4 82152 Planegg, Germany; LMU München, Faculty of Biology, Plant Evolutionary Cell Biology, Großhaderner Str. 2-4 82152 Planegg, Germany; LMU München, Faculty of Biology, Zoology, Großhaderner Str. 2-4 82152 Planegg, Germany

**Keywords:** *Arabidopsis thaliana*, heat acclimation, photosynthesis, subcellular carbohydrate metabolism, invertases, serial block-face scanning electron microscopy

## Abstract

In many plant species, exposure to a changing environmental temperature regime induces an acclimation response which ultimately increases a plant’s thermotolerance. Under elevated temperature, membrane systems need remodelling to counteract de-stabilising thermodynamic effects. This also affects photosynthesis and carbohydrate metabolism due to heat affected protein functions, enzyme activities and transport processes across membrane systems. In the present study, a combination of electrolyte leakage assays and chlorophyll fluorescence measurements was applied to quantify heat tolerance before and after heat acclimation of *Arabidopsis thaliana* at different temperature regimes. Subcellular carbohydrate concentrations were determined in a combined approach of non-aqueous fractionation and 3D reconstruction of mesophyll cells and subcellular compartments using serial block-face scanning electron microscopy. Across temperature regimes between 32 °C and 38 °C, 7 days heat acclimation at 34 °C was found to most efficiently increase tissue heat tolerance. Under such conditions, cytosolic sucrose concentrations were stabilised by a shift of sucrose cleavage rates into the vacuolar compartment while invertase-driven cytosolic sucrose cleavage was found to be efficiently quenched by fructose and glucose acting as competitive and non-competitive inhibitors, respectively. Finally, this study provides strong evidence for a sucrose concentration gradient from the cytosol into the vacuole which might directly affect the physiological role and direction of proton gradient-driven sugar transport across the tonoplast.

## Introduction

Changing temperature regimes have diverse effects on plant growth, development and metabolism. While a sudden and strong temperature drop or increase typically results in irreversible tissue damage and yield loss, a constant and moderate change of temperature induces an acclimation response which increases the temperature tolerance. This acclimation response represents a multigenic process and comprises, among others, signalling cascades, reprogramming of photosynthesis and of the primary and secondary metabolism (Herrmann et al., 2019; Garcia-Molina et al., 2020; Seydel et al., 2022). Increasing temperatures due to global warming have been shown to negatively impact the fitness of plants in their natural habitat and, especially in combination with drought, the yield of crop species (Lippmann et al., 2019; Gampe et al., 2021). Thus, understanding and predicting plant heat response and acclimation capacity is a vital component to understand and deal with the impact of globally increasing temperatures on plants.

Heat is perceived by a changing membrane permeability which, for example, influences transmembrane calcium flux (Ranty et al., 2016; Sajid et al., 2018). Also, reactive oxygen species, kinase and phosphatase activation, phytohormone cascades, activation of transcription factors and heat shock proteins (HSPs) are involved in perception of heat (Iba, 2002; Qu et al., 2013). The immediate recognition of a changing temperature regime is central to stabilise photosynthesis and metabolism. Photosynthesis is known to be a highly temperature sensitive process, being inhibited both at low and high temperatures (Berry and Björkman, 1980). The photosynthetic performance can already be impacted negatively at moderate heat by Rubisco inactivation (Law and Crafts-Brandner, 1999; Crafts-Brandner and Salvucci, 2000; Salvucci and Crafts-Brandner, 2004; Yamori et al., 2014). The thylakoids are influenced by heat-induced changes of membrane properties and the regulatory connection between ATP synthesis and electron transport can be disrupted (Havaux, 1996; Pastenes and Horton, 1996; Bukhov et al., 1999; Bukhov et al., 2000). However, actual damage to photosystem II, quantified by measuring the critical temperature at which the minimal chlorophyll a fluorescence is rapidly increasing, is mostly occurring between 40°C to 55°C, depending on the plant species and growth environment (Terzaghi et al., 1989; Zhu et al., 2018).

Carbohydrates are direct products of photosynthesis and represent energy source and substrates for diverse anabolic pathways. Recent work has shown that carbohydrates are also an important factor in the thermomemory of the shoot apical meristem (Olas et al., 2021). Further, it has been shown that RGS1, a plasma membrane located glucose sensor, is connected to regulation of thermotolerance in tomato, and externally applying glucose to the plants actually increased their thermotolerance (Wang et al., 2024). The enormous plasticity of heat responses in carbohydrate metabolism can also be seen when comparing heat treatments of different duration and severity. Comparing moderate with severe transient heat exposure revealed that sucrose phosphate synthase (SPS) activity negatively correlates with stability of CO_2_ assimilation rates under elevated temperature (Seydel et al., 2022). Another study showed that, in *Arabidopsis*, the amount of primary carbohydrates, e.g., sucrose, raffinose and maltose, increases after treatment with 40°C for up to 240 minutes which represents a shared feature with cold stress (Ahsan et al., 2010). Additionally, within the first 24 hours of heat exposure, 15 % of upregulated proteins in soy bean were found to be related to carbohydrate metabolism, but proteins that were associated with carbon assimilation and photosynthesis were found to be downregulated in the same plants (Ahsan et al., 2010).

Pathways of cellular plant carbohydrate metabolism are located in different subcellular compartments. For example, the Calvin-Benson-Bassham cycle (CBBC) takes place in the chloroplasts while sucrose biosynthesis is catalysed in the cytosol. The high degree of compartmentalisation of plant cells impacts the analysis and understanding of their metabolic pathways profoundly (Lunn, 2007). The method of non-aqueous fractionation (NAF) has been applied to resolve compartment-specific metabolic regulation (Gerhardt and Heldt, 1984; Fürtauer et al., 2016; Hernandez et al., 2023). This method allows for continuous quenching of metabolism and prevents enzymatic interconversion of metabolites after sampling and during fractionation. Correlation of metabolite abundance with marker enzyme activity, or marker protein abundance, provides information about relative metabolite distributions over analysed compartments (Fürtauer et al., 2019). If available, absolute amounts of metabolites can then be multiplied with relative distributions to provide an estimate of compartment-specific absolute metabolite amounts. A current limitation is the estimation of effective subcellular metabolite concentrations which also needs to consider compartment-specific volumes.

In the present study, we have combined leakage assays and chlorophyll fluorescence measurements with the NAF methodology and serial block-face scanning electron microscopy to quantify thermotolerance and photosynthetic efficiency together with carbohydrate concentrations in the plastids, cytosol and vacuole of leaf mesophyll cells of *Arabidopsis thaliana*. Effective compartment-specific concentrations were determined before and after acclimation to 34 °C to unravel heat-induced regulation of subcellular carbohydrate metabolism.

## Methods

### Plant material

Plants of *Arabidopsis thaliana,* Columbia-0 (Col-0), were grown at 22 °C in the greenhouse (∼100-125 µmol m^-2^ sec^-1^, ∼12 h/12 h light/dark). After 4 weeks of growth, at initial bolting stage, the control plants were harvested. For heat acclimation, plants were transferred to the respective temperature at a light intensity of 100 µmol m^-2^ sec^-1^, 12 h/12 h light/dark and watered regularly to prevent drought stress. Temperature treatment included 32 °C, 34 °C, 36 °C, and 38 °C. After 7 days of heat treatment, the plants were harvested. For 38 °C, harvesting took place after 3 days of heat treatment due to high mortality of the plants beyond that timepoint. Harvesting for analysis of metabolism took place at mid-day, i.e., after 6 h in light, by cutting the plants at the hypocotyl and plunge-freezing them in liquid nitrogen. Frozen plant material was stored at −80 °C until it was ground to a fine powder and freeze dried. Samples for electron microscopy were collected at the end of the night to prevent starch accumulation in the chloroplasts.

### Electrolyte leakage

An electrolyte leakage assay was performed on intact leaves from control plants and acclimated plants. Two leaves per sample were cut from one plant and fully submerged in 7 ml of distilled H_2_O. The samples were heated to 46°C for 0, 30, 45, and 60 minutes. They were cooled down and shaken over night at room temperature. Then, 1 ml of water was taken from the samples, diluted 1:4 in H_2_O and initial electrical conductivity EC_initial_ [µS cm^-1^] was determined. Then, the samples were heated to 95 °C for 1 h and total electrical conductivity EC_total_ [µS cm^-1^] was determined again. The index of injury I_d_ was calculated individually for the different timepoints with the fractional release of electrolytes from non-heated and heated samples, R_0_ and R_t_, (**Equations 1-3**; Flint et al. (1967)).

*Equation 1: Index of Injury Id from exposure to temperature*

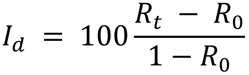

*Equation 2: Fractional release of electrolytes from non-heated sample R0 (timepoint 0 minutes)*

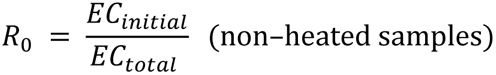

*Equation 3: Fractional release of total electrolytes from heated sample Rt (timepoints 30, 45, 60 and 75 minutes)*

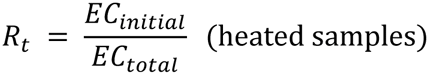

### Chlorophyll fluorescence

To evaluate the effect of prolonged heat exposure on photosystem II, maximum quantum yield (F_v_/F_m_) was recorded by supplying a saturating light pulse after 15 minutes of dark adaptation at 22 °C (WALZ JUNIOR-PAM®; Heinz Walz GmbH, Germany).

### Metabolite quantification

Amounts of carbohydrates were determined as described before (Kitashova et al., 2023). Soluble carbohydrates were extracted twice with 400 µl 80 % ethanol at 80 °C for 30 minutes. The supernatants were combined and dried for sugar analysis. The pellet was used for starch quantification by amyloglucosidase digestion and photometric detection of glucose equivalents by a coupled glucose oxidase/peroxidase/o-dianisidine reaction. Sucrose content was determined by an anthrone assay, whereas glucose concentration was determined by a coupled hexokinase/glucose-6-phosphate dehydrogenase assay, utilizing absorption measurement of produced NADPH + H^+^. Fructose quantification followed glucose quantification by addition of PGI to the reaction buffer.

### Invertase and SPS activity measurements

Activities of vacuolar (acidic), cytosolic (neutral) and cell wall-bound invertases as well as of sucrose phosphate synthase (SPS) were quantified as described earlier with slight modifications (Nägele et al., 2012; Kitashova et al., 2022). Invertases were extracted on ice in extraction buffer (50 mM HEPES-KOH, pH 7.5, 5 mM MgCl_2_, 2 mM EDTA, 1 mM phenylmethylsulfonylfluoride, 1 mM DTT, 0.1 % (v/v) Triton-X-100, 10 % (v/v) glycerol). After centrifugation, the supernatant was analysed for vacuolar and cytosolic invertase, whereas the pellet was resuspended in extraction buffer to analyse the cell wall-bound invertase activity. After incubation of the supernatant at 30 °C in acidic reaction buffer (20 mM sodium acetate, pH 4.7, 100 mM sucrose) for vacuolar and cell wall invertase measurements, or neutral reaction buffer (20 mM HEPES-KOH, pH 7.5, 100 mM sucrose) for cytosolic invertase, the solution was neutralised with 1 M NaH_2_PO_4_, heated to 95 °C to stop the enzymatic reaction, and centrifuged. The glucose content in the supernatant was quantified photometrically by a coupled glucose oxidase/peroxidase/o-dianisidine reaction.

SPS was extracted in extraction buffer (50 mM HEPES-KOH, pH 7.5, 20 mM MgCl_2_, 1 mM EDTA, 2,7 mM DTT, 10 % (v/v) glycerol, 0.1 % (v/v) Triton-X-100) on ice and centrifuged. The supernatant was incubated with reaction buffer (50 mM HEPES-KOH, pH 7.5, 15 mM MgCl_2_, 3 mM DTT, 35 mM UDP-glucose, 35 mM fructose-6-phosphate, 140 mM glucose-6-phosphate) at 25 °C and the reaction was stopped by adding 30 % KOH and heating the solution to 95 °C. Sucrose was quantified photometrically with an anthrone assay.

### Nonaqueous fractionation

Nonaqueous fractionation (NAF) was performed as described previously (Fürtauer et al., 2016; Hernandez et al., 2023). Briefly, 10-15 mg of lyophilised plant material was suspended in 1 ml of a heptane (C_7_H_16_; “7H”) -tetrachlorethylene (C_2_Cl_4_; “TCE”) mixture with a density of *ρ*=1.35 g cm^-3^ and sonicated on ice in 30 sec pulses with pauses of 1 min for a total of 20 minutes (Hielscher UP200St Ultrasonic Homogenizer, 170 W, 100 % power setting; Hielscher Ultrasonics GmbH, Teltow, Germany). The sonicated material was centrifuged for 20 min at 4°C and 20000 x g. The supernatant was stored on ice and the pellet was suspended in a 7H-TCE mixture of higher density and sonicated for 10 sec to facilitate dissolving of the pellet. Subsequently, the material was centrifuged, and the new pellet suspended in higher density 7H-TCE mixture. This process was repeated with mixtures of increasing density, ranging from *ρ* =1.35 g cm^-3^ to *ρ* =1.6 g cm^-3^. The fractions were split equally and dried in a vacuum desiccator. Pellets were stored at −20 °C until subsequent analysis. One aliquot per sample was used for photometric marker enzyme measurements for vacuole, cytosol and chloroplast and the other for photometric evaluation of sugar content.

### Marker enzyme activities

For photometric measurement of marker enzyme activities, the dried pellets of fractions were suspended in 750 µl extraction buffer (50 mM Tris-HCl, pH 7.3, 5 mM MgCl_2_, 1 mM DTT), incubated on ice for 10 min and centrifuged at 4°C and 20,000 g for 10 min. The supernatant was used for enzyme activity quantification. Plastidial pyrophosphatase was used as marker enzyme for the chloroplast, cytosolic uridine diphosphate (UDP) glucose pyrophosphorylase (UGPase) for the cytosol and vacuolar acidic phosphatase for the vacuole (Fürtauer et al., 2019). Plastidial pyrophosphatase was assayed as described earlier (Jelitto et al., 1992) and inorganic phosphate detected by molybdenum blue reaction (Murphy and Riley, 1962). UGPase was quantified photometrically as described before (Zrenner et al., 1993). Acidic phosphatase from the vacuole was measured according to Boller and Kende (1979), with some modifications. The assay buffer consisted of 125 mM sodium acetate and 0.125 % Triton-X 100 and was adjusted to pH 4.8 with acetic acid, whereas the substrate for the detection was composed of 1 mg/ml 4-nitrophenylphosphate in assay buffer.

### Sample fixation for microscopy

Several leaves per plant were fixed for microscopy. The plants were harvested at the end of the dark period to minimise starch content in the plastids, which improves visibility of thylakoid membranes. The leaves were cut into 1 mm^2^ pieces in fixation buffer (75 mM cacodylate, 2 mM MgCl_2_, pH 7.0) supplemented with 2.5 % glutaraldehyde and stored at 4°C for several days until further processing.

### Light and transmission electron microscopy

For light and transmission electron microscopy, fixation was carried out as described before (Garcia-Molina et al., 2021). After post-fixation with 1 % (w/v) OsO_4_, the samples were contrasted *en bloc* with 1 % (w/v) uranyl acetate in 20 % acetone, dehydrated with a graded acetone series and embedded in Spurr’s resin of medium rigidity (Spurr, 1969). For TEM, ultrathin sections of approximately 60 nm were contrasted with lead citrate (Reynolds, 1963) and examined with a Zeiss EM 912 transmission electron microscope with an integrated OMEGA energy filter, operated at 80 kV in the zero-loss mode (Carl Zeiss AG, Oberkochen, Germany). Images were acquired with a 2k x 2k slow-scan CCD camera (TRS Tröndle Restlichtverstärkersysteme, Moorenweis, Germany). For light microscopy, semi thin sections of 1µm were examined with a Zeiss Axiophot microscope and a SPOT Insight camera.

### Serial block-face scanning electron microscopy

For serial block-face scanning electron microscopy (SBF-SEM), 1 mm^2^ pieces of *A. thaliana* leaves were fixed and stained following a protocol based on Hua et al. (2015). In brief, with intermittent washing steps, the samples were fixed as described above, treated with 2 % OsO_4_ + 1.5 % potassium ferrocyanide on ice, with thiocarbohydracid solution at RT, again 2 % OsO_4_, 1 % uranyl acetate, then lead aspartate at 60 °C. Following an ascending ethanol and acetone series, the samples were embedded in epon resin hard 812, mounted on aluminium stubs with conductive glue, trimmed to 500 µm cubes and sputtered with 20 nm gold.

Serial sectioning and imaging took place on a ThermoFisher Apreo VS blockface-scanning electron microscope (Thermo Fisher Scientific Inc., Waltham, USA) in low vacuum at 2.1 kV, 100-200 pA and a pixel dwell time of 3 µs. Digital image stacks (greyscale, 8 bit) of 8192 x 8192 pixels at 20 nm pixel size and 40 nm cutting thickness were generated and post-aligned with Fiji, with stack 1 amounting to 1872 planes and stack 2 amounting to 1450 planes.

### Segmentation of cellular compartments

The SBF-SEM image stacks were processed with the Amira Pro software (Versions 2019-2024.1, Thermo Fisher Scientific). In a fraction of the two datasets, chloroplasts, vacuole, cytoplasm and nucleus were segmented manually with a graphic tablet. Smaller organelles such as mitochondria, peroxisomes, Golgi apparatus and endoplasmic reticulum were not segmented, but were included in the cytoplasm material. The Python Deep Learning environment of Amira was utilised for automated segmentation of the datasets with the manually segmented data serving as ground truth. Learning settings were adjusted between training steps and are summarised in the supplements (**Supplementary Table ST1**). The resulting labels were checked for mislabelling and corrected manually.

### Data analysis and calculation of subcellular volumes

Subcellular sugar content was correlated to the marker enzyme measurements with the “NAFalyzer” app to estimate subcellular sugar concentrations relative to sample dry weight (Hernandez et al., 2023).

Leaf discs of 4 mm radius were punched out and subsequently dried to quantify fresh and dry weight (n = 45). Average leaf height (n ≥ 22) from light micrographs was multiplied with disc size to determine the leaf disc volume. The volume to dry weight ratio was calculated and corrected for gas space percentage of the segmented SBF-SEM dataset. Volume per dry weight ratio of plastids, cytosol and vacuole were calculated from the percentages of the compartments from the SBF-SEM dataset and further combined with the subcellular sugar amount to calculate the absolute subcellular sugar concentrations in mM (**Supplementary Tables ST2 and ST3**). The percentage of cell types in the leaf were measured in light microscopic images of semi thin sections of embedded leaf material (n = 4). Image analysis of electron and light micrographs was carried out with Fiji. Data analysis and statistics were carried out with R (The R Project for Statistical Computing; https://www.r-project.org) and Microsoft Excel (https://www.microsoft.com). The development of the R shiny app for simulation of Michaelis-Menten kinetics was supported by ChatGPT (October 2024). The code is provided via GitHub: https://github.com/cellbiomaths/Shiny_MM_simulation.

## Results

### Electrolyte leakage indicates efficient Arabidopsis heat acclimation between 32 °C and 34 °C

Susceptibility to heat stress was quantified by quantifying the electrolyte leakage of leaf tissue to estimate membrane damage, described by the Index of Injury *I*_d_. Leaf tissue acclimated to 22 °C was damaged to ∼80 % after 45-60 min of incubation in 45 °C water (**Fig. 1 A**). Heat acclimation of leaf tissue after 7 days was most efficient for acclimation temperatures 32 °C and 34 °C which resulted in significant reduction of *I*_d_ values (ANOVA, **Fig. 1 B, C**). Acclimation at 36 °C (7 days) and 38 °C (3 days) resulted in a slight decrease of *I_d_* when compared to 22 °C plants, but the condition effect was not significant (**Fig. 1 D, E**). This was also in accordance with the growth phenotype of the plants after the heat treatment which indicated severe tissue damage at 36 °C and 38 °C, whereas for 32 °C and 34 °C, only an early induction of the inflorescence and slight yellowing of the leaves was observed (**Supplementary Figure SF1**).

**Figure 1.**
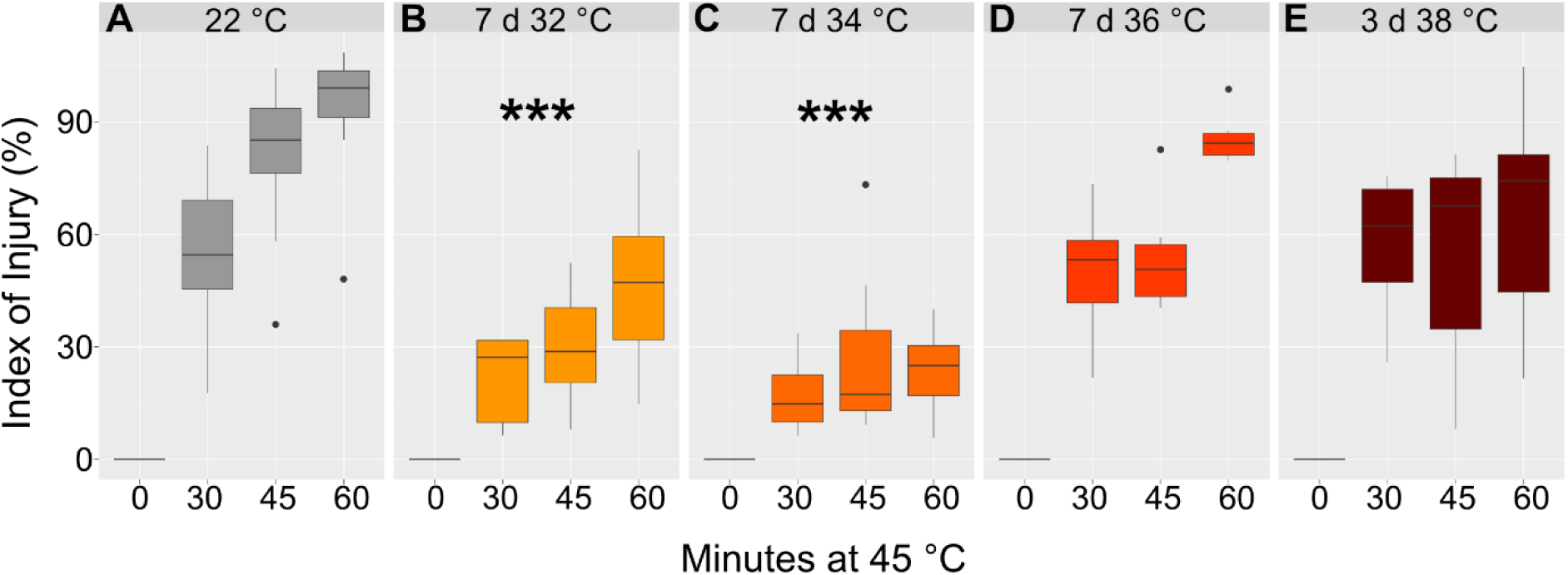
Heat tolerance of leaf tissue before and after acclimation. The Index of injury [%] was based on quantified electrolyte leakage quantification of leaf tissue of plants grown at 22°C (**A**) and heat acclimated (**B**) 7 days at 32°C, (**C**) 7 days 34°C, (**D**) 7 days 36°C, and (**E**) 3 days 38°C. Asterisks indicate significant difference from control (ANOVA and Tukey HSD post-hoc test, *** p < 0.001), n=6.

To reveal how elevated temperature affected photosynthetic efficiency, the maximum quantum yield of PSII (F_v_/F_m_) was quantified (**Fig. 2**). Under control conditions (22 °C), F_v_/F_m_ values were >0.8 while they dropped significantly after 7 days at 32 °C, 34 °C and 36 °C to values between 0.76 and 0.78. After 3 days at 38 °C, F_v_/F_m_ showed the strongest decrease to a median of ∼0.64, and the data variance distinctly increased (**Fig. 2**, dark red box).

**Figure 2.**
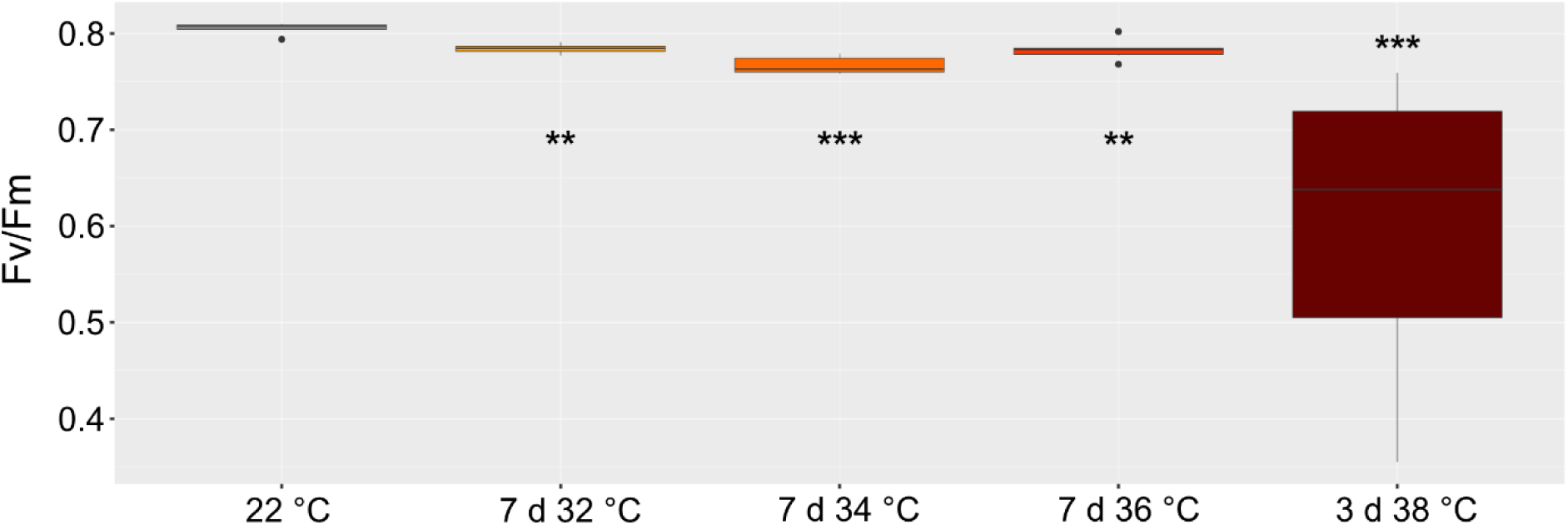
Maximum quantum yield of photosystem II (F_v_/F_m_) as a function of acclimation duration and temperature. Colours indicate acclimation temperature: grey: 22 °C; yellow: 7 days 32 °C, orange: 7 days 34 °C, light red: 7 days 36 °C, dark red: 3 days 38 °C. Asterisks indicate significant difference from control (ANOVA and Tukey HSD post-hoc test, ** p < 0.01, *** p < 0.001), n = 6.

### Heat response of the central carbohydrate metabolism

Starch amounts decreased significantly under elevated temperatures (**Fig. 3 A**). Plants acclimated for 7 days at 32 °C had approximately 20 % of starch found in tissue of non-acclimated plants while amounts dropped even more for higher temperature regimes.

**Figure 3.**
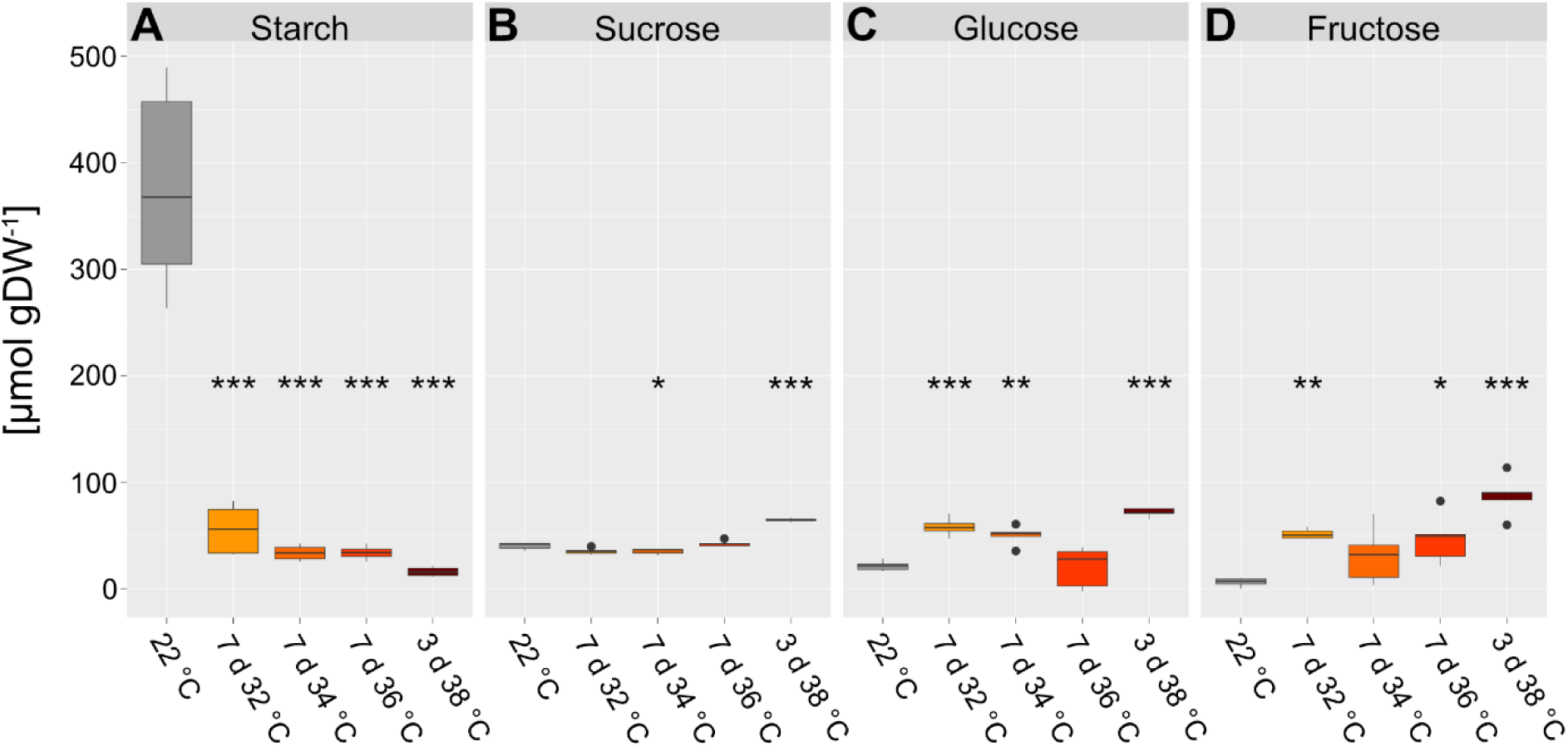
Metabolite amount per gram dry weight in control and acclimated plants (grey: 22 °C, yellow: 32 °C, orange: 34 °C, light red: 36 °C, dark red: 38 °C). (**A**) Starch amount in C6 equivalents. (**B**) Sucrose amount. (**C**) Glucose amount. (**D**) Fructose amount. Asterisks indicate significant difference from control (ANOVA and Tukey HSD post-hoc test, * p < 0.05, ** p < 0.01, *** p < 0.001), n≥5.

While, compared to non-acclimated plants, sucrose amounts also dropped in plants acclimated at 34 °C, they significantly increased approximately 1.5-fold after 3 days at 38 °C (**Fig. 3 B**). For both glucose and fructose amounts were found to significantly increase during heat exposure with an exception at 36 °C for glucose and 34 °C for fructose (**Fig. 3 C, D**).

### Dynamics of subcellular sugar compartmentation under heat

Based on the finding that heat acclimation resulted in highest tolerance after 7 days at 34 °C (**Fig. 1 A, C**), this condition was chosen for further detailed analysis of subcellular compartmentation of carbohydrates. The compartment-specific relative distribution of soluble sugars sucrose, glucose and fructose was quantified before (22 °C) and after heat acclimation (7d 34 °C).

For subcellular sucrose distribution it was observed that heat induced a significant shift from chloroplasts to the vacuole (**Fig. 4 A**). Due to heat exposure, the plastidial sucrose proportion decreased to less than 10 % while it increased to ∼80 % in the vacuole. This shift was neither observed for glucose nor for fructose (**Fig. 4 B, C**). Under heat, glucose was slightly shifted from the vacuole into the cytosol, and fructose proportions remained relatively constant.

**Figure 4.**
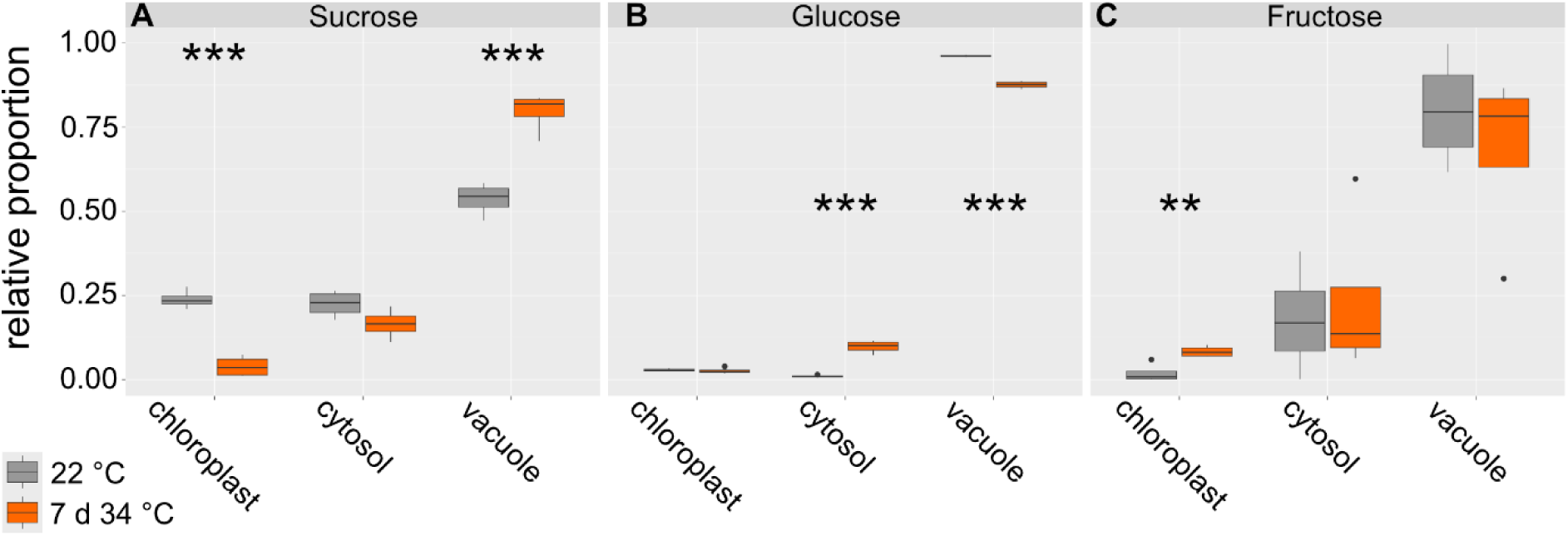
Effects of heat acclimation on subcellular distribution of soluble carbohydrates. Relative proportions for (**A**) sucrose, (**B**) glucose and (**C**) fructose were resolved for chloroplasts, cytosol and vacuole before (22 °C; grey boxes) and after heat acclimation at 7d 34 °C (orange boxes). Asterisks indicate significance between both conditions (ANOVA and Tukey HSD post-hoc test; ** p < 0.01, *** p < 0.001), n = 4.

The observed shifts in the relative proportion of sugars indicated a heat-induced regulation of subcellular compartmentation of carbohydrate metabolism. To reveal the effect of these relative shifts on compartment-specific metabolite concentrations, volumes of chloroplasts, cytosol and vacuole were experimentally resolved by three-dimensional SBF-SEM for mesophyll tissue at 22 °C and 34 °C (**Fig. 5; Supplementary Figure SF2**).

**Figure 5.**
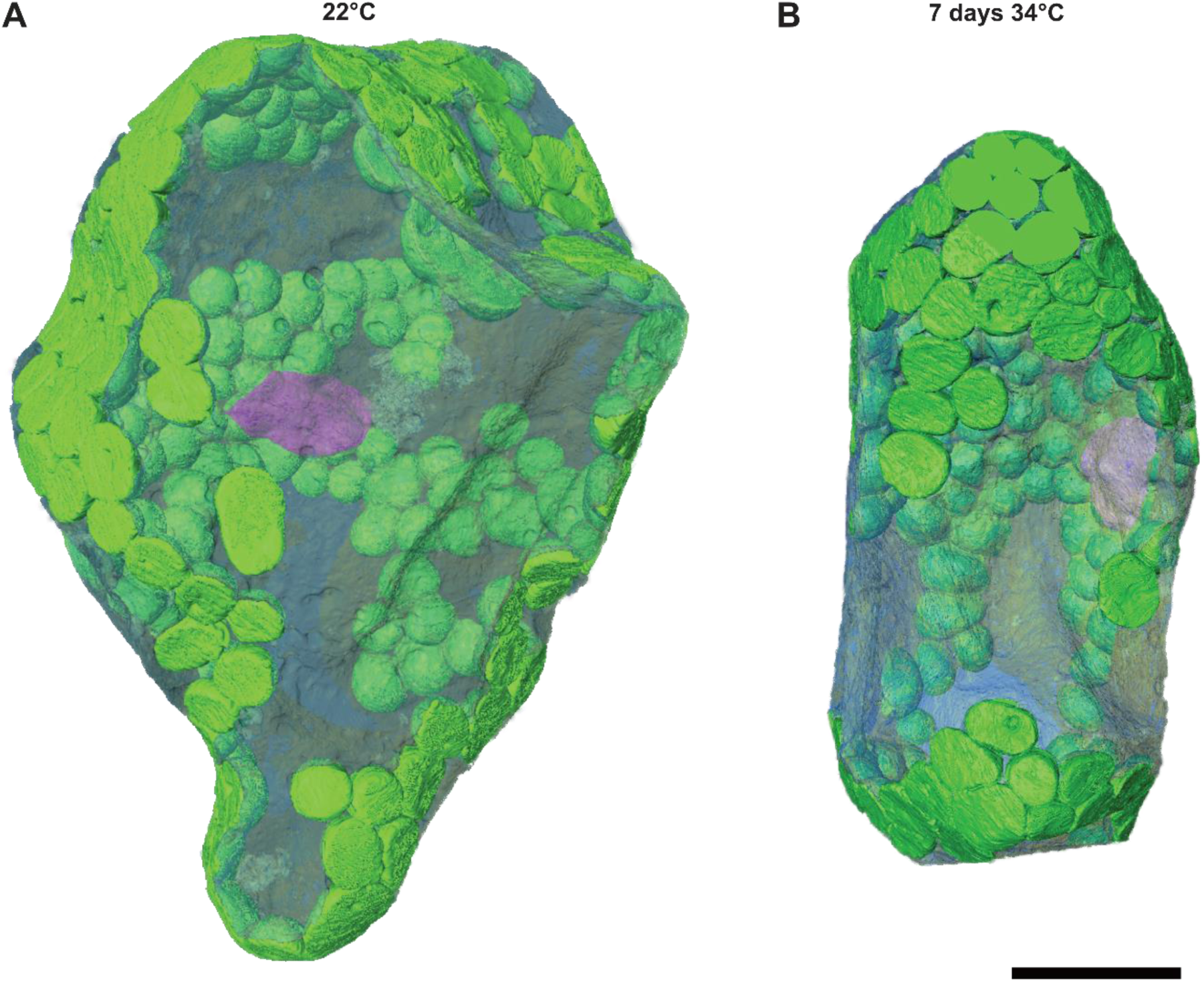
3D models of mesophyll cells. (**A**) Control (22 °C) spongy mesophyll cell. (**B**) Heat treated (7 days 34 °C) palisade mesophyll cell. Chloroplasts are shown in green, the nucleus in pink and the vacuole in translucent blue. The cytosol encompassing other organelles such as mitochondria, Golgi apparatus or endoplasmic reticulum, surrounding the vacuole and the chloroplasts is omitted in this view. Scalebar: 20µm.

Evaluation of compartmental proportions revealed ∼14 % chloroplasts, 3.5 % cytosol and ∼83 % vacuole at 22 °C and 34 °C (**Table I**). Together with the measured dry weight to fresh weight ratio and the average thickness of a leaf, fractions of compartment volumes in fresh leaves were determined. At 22 °C, the tissue volume per gram dry weight was about 1.5-fold higher than in heat treated plants. Considering the estimated gas space from SBF-SEM analysis, this finally allowed for estimating the volume of cell material which was 10,932 mm³ gDW^-1^ at 22 °C and 8,267 mm³ gDW^-1^ at 34 °C. This information was combined with the compartmental proportions to estimate volumes of chloroplasts, cytosol and vacuole with the dimension of ml gDW^-1^ (**Table I**).

**Table I.** Compartment volumes and leaf measures. Proportions in % were determined from SBF-SEM data, leaf height from light micrographs. Leaf disc volume was calculated from leaf height and punchout size (4mm radius). Dry weight per leaf disc was determined by drying and weighing leaf discs. Volume per gDW was calculated from leaf disc volume and dry weight. Gas space percentage was acquired from SBF-SEM data and used to determine the volume of cell material per gDW. This volume and the compartment proportions were used to estimate the volume of the different cell compartments per gDW. Detailed calculations are collected in the supplements (**Supplementary Table ST2**).

| Measurement | Control | 34°C |
| --- | --- | --- |
| Proportion chloroplasts | 13.89 % | 13.65 % |
| Proportion cytosol | 3.51 % | 3.63 % |
| Proportion vacuole | 82.6 % | 82.72 % |
| Leaf height [mm] | 0.187 | 0.138 |
| Leaf disc volume [mm<sup>3</sup>] | 9.415 | 6.916 |
| Dry weight per leaf disc [mg] | 0.622 | 0.645 |
| Volume per gDW [mm<sup>3</sup> g<sup>-1</sup>] | 15,131.2 | 10,731.9 |
| Percentage gas space (SBF-SEM) | 0.278 | 0.229 |
| Volume gas space [mm<sup>3</sup> gDW<sup>-1</sup>] | 4,199.5 | 2,465.5 |
| Volume cell material [mm<sup>3</sup> gDW<sup>-1</sup>] | 10,931.8 | 8,266.5 |
| Estimated volume chloroplasts | 1.52 ml gDW<sup>-1</sup> | 1.13 ml gDW<sup>-1</sup> |
| Estimated volume cytosol | 0.38 ml gDW<sup>-1</sup> | 0.30 ml gDW<sup>-1</sup> |
| Estimated volume vacuole | 9.03 ml gDW<sup>-1</sup> | 6.84 ml gDW<sup>-1</sup> |

This revealed that, although relative proportions of analysed compartments remained stable under heat, absolute compartment volumes decreased by about 25 % after 7 d at 34 °C due to a reduced volume of leaf tissue per gDW.

The percentage of different cell types in a sampled leaf was then determined by light microscopy (**Table II**). This analysis revealed approximately 57 % mesophyll cells, 15 % epidermal cells, 2 % vascular bundle cells and 26 % gas space at 22 °C. During heat, mesophyll and epidermal cell proportions slightly increased to 58 % and 16 %, respectively. Vascular bundles did not change, whereas the gas space decreased to 24 %.

**Table II:**
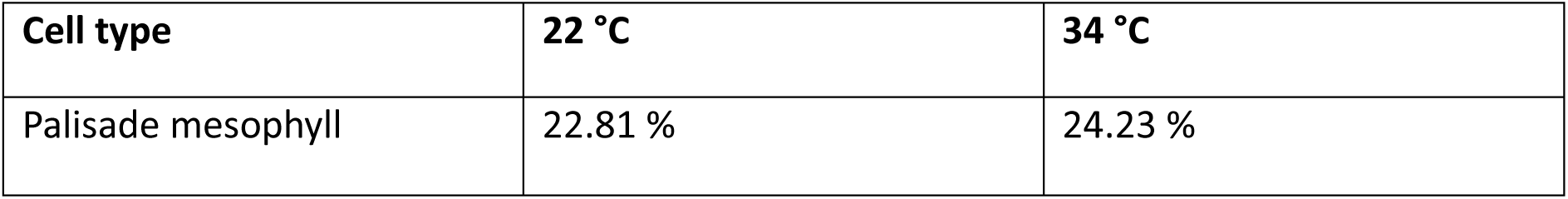
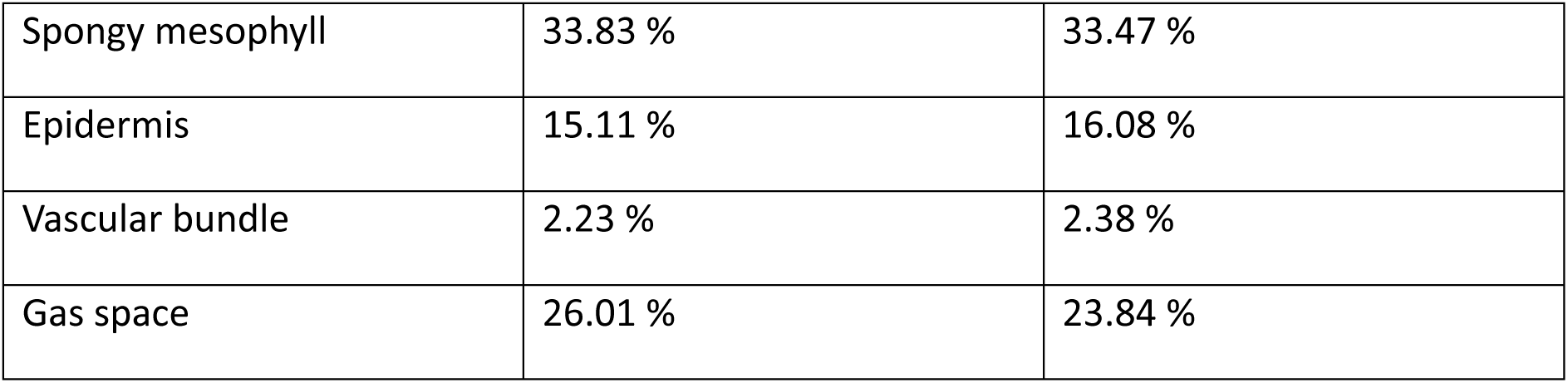
Percentage of cell types in a leaf section. The area of cell types was measured in light micrographs of semi-thin sections of embedded leaf material, n = 4

| Cell type | 22 °C | 34 °C |
| --- | --- | --- |
| Palisade mesophyll | 22.81 % | 24.23 % |
| Spongy mesophyll | 33.83 % | 33.47 % |
| Epidermis | 15.11 % | 16.08 % |
| Vascular bundle | 2.23 % | 2.38 % |
| Gas space | 26.01 % | 23.84 % |

Combining the absolute sugar amounts with NAF-derived subcellular proportions revealed absolute sugar amounts of chloroplasts, cytosol and vacuole at 22 °C and after 7 d at 34 °C (**Fig. 6 A-C**). Absolute sucrose amount differed significantly between both conditions across all compartments (**Fig. 6 A**). The trend observed for relative proportions (see **Fig. 4**) was augmented, and cytosolic amounts now also differed significantly. Absolute amounts of glucose were significantly elevated due to heat acclimation across all compartments (Fig. 6 B). In the vacuole, this contrasted the relative proportions which significantly decreased at 34 °C (compare **Fig. 4 B**). However, due to a higher total amount of glucose under heat, this relative decrease still resulted in an absolute increase in the vacuolar compartment. Fructose was found to significantly accumulate in the vacuole which was not observed for relative proportions (compare **Fig. 4 C**). As described for glucose, this was due to an increase in total fructose amounts. Yet, fructose dynamics were less significant than for glucose and sucrose.

**Figure 6.**
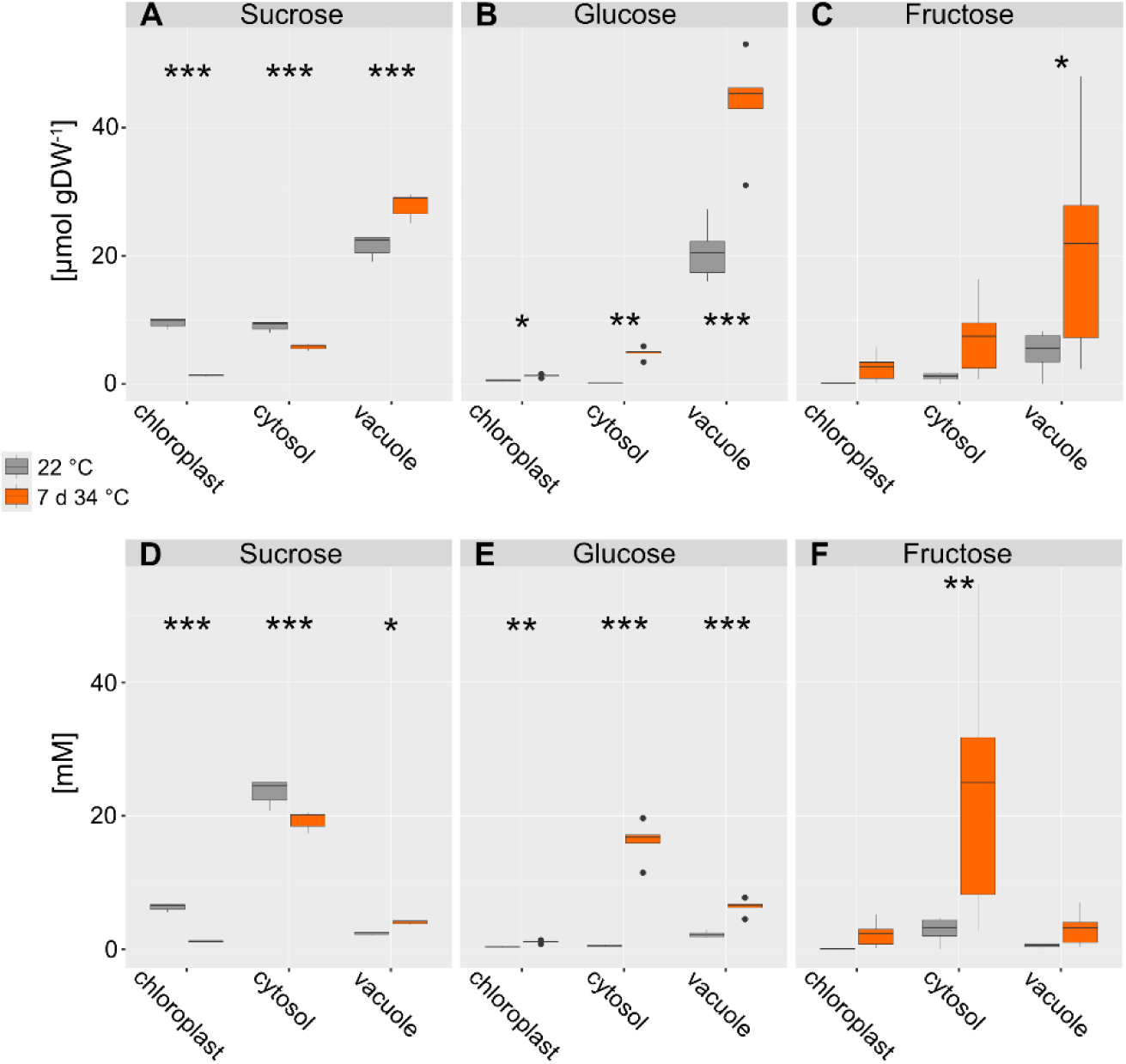
Compartment-specific sugar amounts and concentrations. Absolute amounts of sugars in compartments chloroplasts, cytosol and vacuole are shown in the upper panel (**A – C**), concentrations in [mM] are shown in the lower panel (**D – F**). Grey boxes: 22 °C, orange boxes: 7d 34 °C. Asterisks indicate significance (ANOVA and Tukey HSD post-hoc test, * p < 0.05, ** p < 0.01, *** p < 0.001), n = 5.

Next, absolute compartment-specific sugar concentrations were derived from amounts normalised to estimated volumes of a mesophyll cell (**Fig. 6 D-F**). Interestingly, this emphasised accumulation effects in the cytosol which now also became significant for fructose (**Fig. 6 F**). In general, while subcellular sugar amounts (in μmol gDW^-1^) were highest in the vacuole, subcellular sugar concentrations (in mM) peaked in the cytosol which was due to the strong discrepancy of vacuolar and cytosolic volumes (see **Table I**).

### Heat-induced dynamics of enzyme activities in sucrose metabolism

Based on the observation that heat significantly affected compartmentation of sucrose and its hydrolytic cleavage products glucose and fructose, activities of central enzyme activities of sucrose biosynthesis (sucrose phosphate synthase; SPS) and cleavage (neutral, acidic and cell wall-associated invertases; nInv, aInv, cwInv) were quantified before and after 7 d at 34 °C (**Fig. 7**). Activities were normalised to the volume of leaf cells, i.e., 10.93 ml gDW^-1^ at 22 °C and 8.26 ml gDW^-1^ at 34 °C. Activities of SPS and nInv decreased significantly under heat (**Fig. 7 A, B**) while cwInv and aInv activities increased non significantly (**Fig. 7 C, D**).

**Figure 7.**
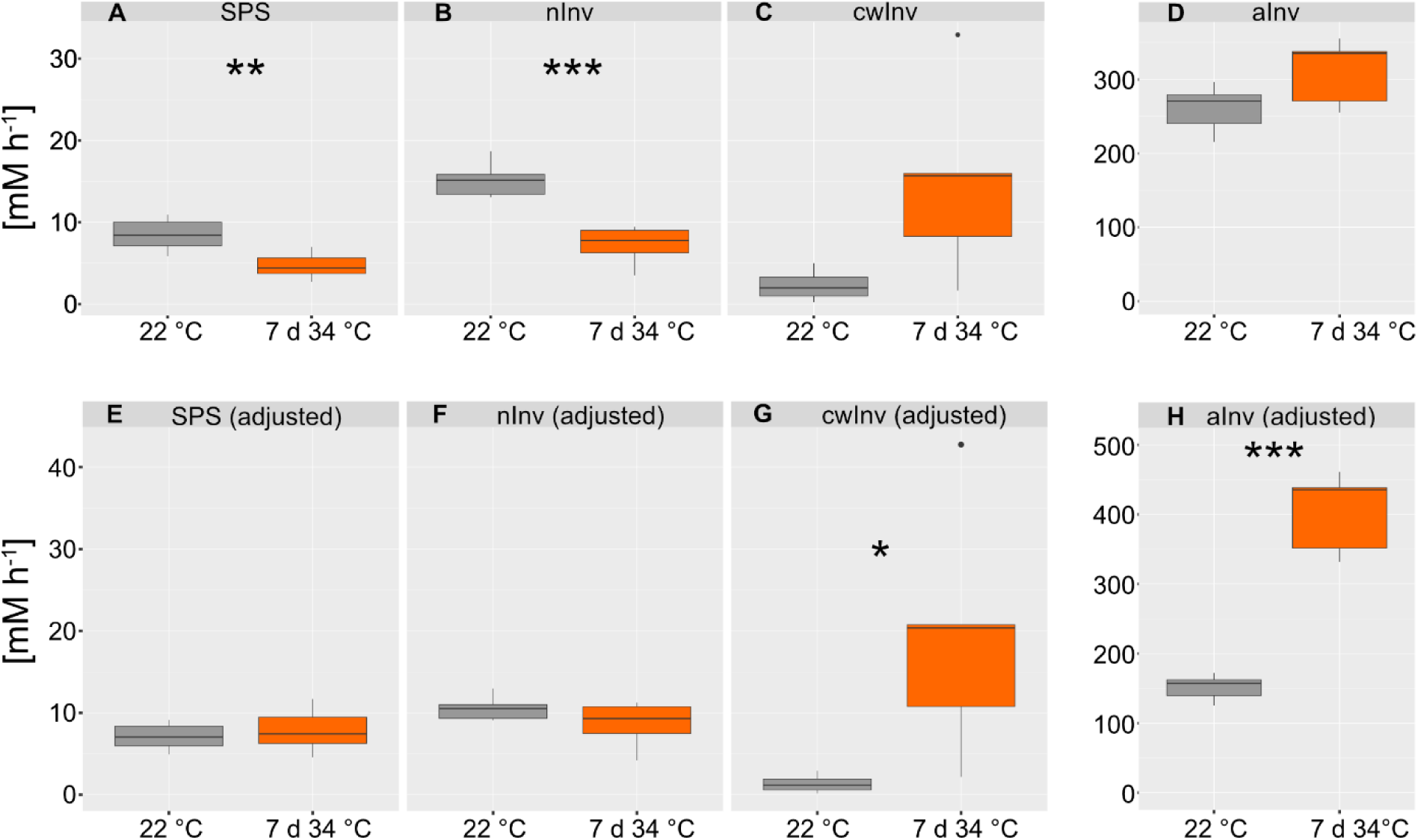
Activities of enzymes in sucrose metabolism. Invertase and SPS activity of plants grown at 22 °C (grey) and 34 °C acclimated plants (orange) were determined under substrate saturation (i.e., V_max_). The upper panel (**A – D**) represents experimentally quantified activities at temperature optimum (SPS: 25 °C; invertases: 30 °C). The lower panel (**E – H**) shows activities which were adjusted to the growth temperatures using the Arrhenius equation (see main text). Asterisks indicate significance in Student’s t-test (* p < 0.05, ** p < 0.01, *** p < 0.001), n = 5.

Enzyme activities were quantified under substrate saturation at temperature optimum, which was 25 °C for SPS and 30 °C for invertases, respectively. To adjust those activities to the growth temperature of the plants, activation enthalpies were applied and used to solve the Arrhenius equation ((Arrhenius, 1889), **Eq. 4**).

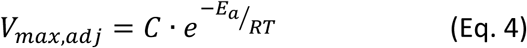

Here, C represents the Arrhenius factor, E_a_ is the activation energy, R is the gas constant and T the temperature. The Arrhenius factor was estimated from experimentally determined enzyme activities at their temperature optimum as described earlier ((Weiszmann et al., 2018); **Supplementary Table ST4**). The adjustment to physiologically more relevant temperature regimes resulted in similar activities of SPS and nInv at 22 °C and 34 °C, respectively (**Fig. 7 E, F**). In contrast, adjustment resulted in significantly different activities of cwInv and aInv at 22 °C and 34 °C (**Fig. 7 G, H**). In summary, thermodynamic adjustment of enzyme activities revealed that both reactions located in the cytosol, SPS and nInv, were efficiently stabilised under heat to maintain similar maximum enzyme activities as under 22 °C. In contrast, both enzymes located in acidic environments, i.e., apoplast (cwInv) and vacuole (aInv), significantly increased in their maximum activities.

Using the subcellular concentrations of sucrose, glucose and fructose together with adjusted activities of enzymes, *in vivo* rates of cytosolic (nInv) and vacuolar (aInv) sucrose cleavage were estimated assuming Michaelis-Menten kinetics with competitive (Frc) and non-competitive (Glc) inhibition ((Sturm, 1999), **Fig. 8**). Simulations showed that, under heat, cytosolic rates of sucrose cleavage were dramatically reduced due to increased cytosolic hexose feedback inhibition of nInv (**Fig. 8 B**). For aInv, which represents soluble acidic invertases with vacuolar localization, V_max_ was significantly increased at 34 °C which resulted in a maintenance of sucrose cleavage rates (**Fig. 8 C, D**) although also vacuolar concentrations of Frc and Glc increased under these conditions (see **Fig. 6**).

**Figure 8.**
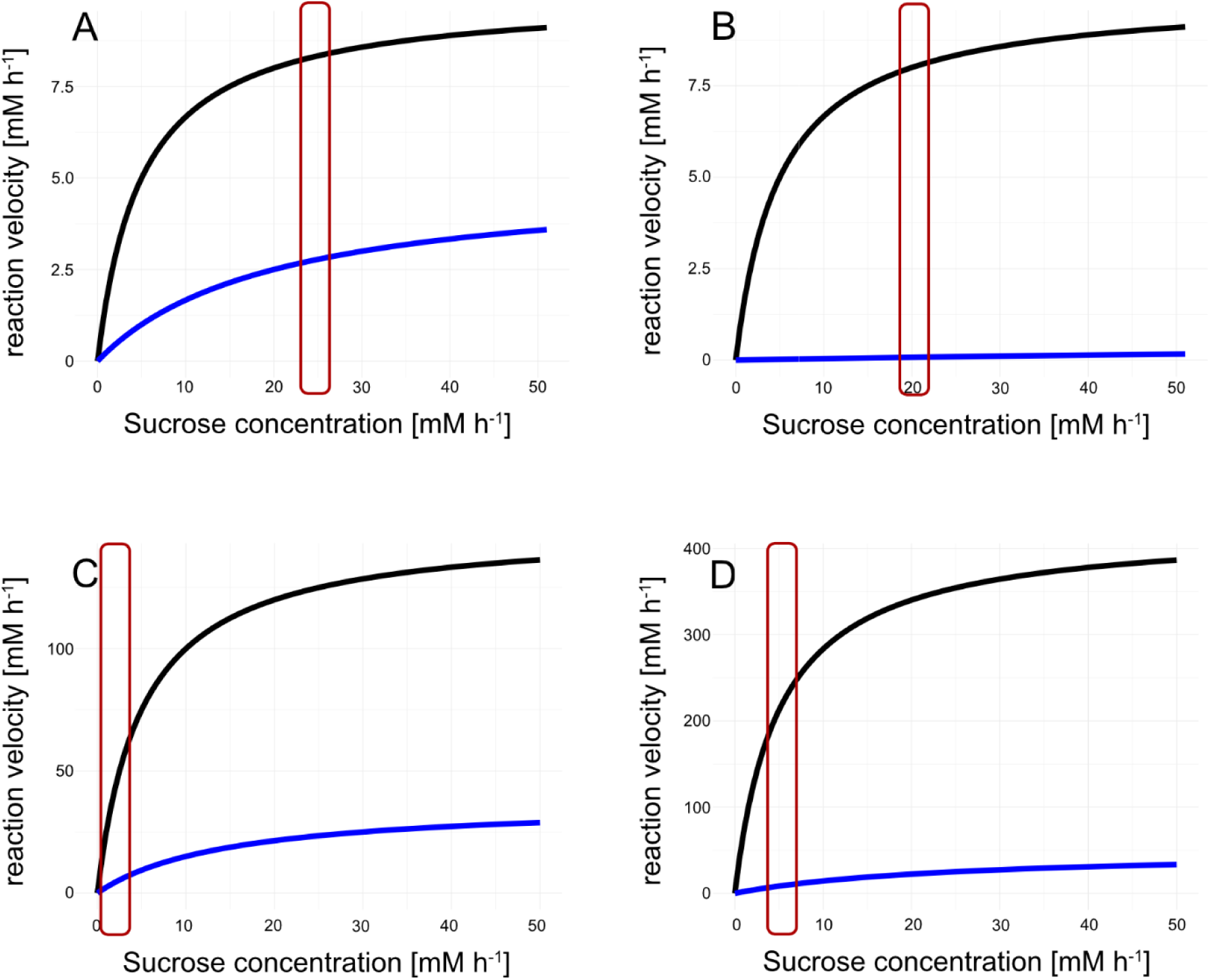
Simulation of Michaelis-Menten kinetics using subcellular metabolite concentrations and temperature-adjusted enzyme activities. **(A)** Simulation of nInv catalysed reaction at 22 °C, **(B)** Simulation of nInv catalysed reaction at 34 °C, **(C)** Simulation of aInv catalysed reaction at 22 °C, **(D)** Simulation of aInv catalysed reaction at 34 °C. black lines: no inhibition; blue lines: combined competitive and non-competitive inhibition by Frc and Glc, respectively. Red boxes indicate the relevant ranges of sucrose concentrations *in vivo*. Parameter settings are provided in the supplement (**Supplementary Table ST5**). For simulations, the code for an R Shiny app is provided on GitHub: https://github.com/cellbiomaths/Shiny_MM_simulation).

## Discussion

A changing temperature regime needs to be efficiently perceived and sensed by plants in order to initiate stress and acclimation response. Particularly, if temperature decreases or rises below or above critical values, molecular and physiological adjustments become essential to prevent irreversible cell and tissue damage. Adjustment and stabilization of photosynthesis and carbohydrate metabolism are central to temperature acclimation because a deflection of involved processes and pathways directly affects plant performance (Anderson et al., 1995; Qu et al., 2023). To quantify heat tolerance of *Arabidopsis thaliana* on levels of tissue structure and photosynthetic efficiency, electrolyte leakage assays were combined with measurements of F_v_/F_m_ in the present study. While a 7-days acclimation period at 32 °C and 34 °C resulted in a significant reduction of leakage of leaf tissue when compared to non-acclimated plants, exposure to higher temperatures did not significantly improve tissue heat tolerance. Also, although F_v_/F_m_ significantly dropped under all tested regimes of elevated temperature, it stabilised at ∼0.75 (except for the 38 °C treatment) which still indicated a relatively high efficiency of PSII. However, these PAM measurements were conducted on green chlorophyll containing tissue while growth phenotypes at 36 °C already showed pale areas (see **Supplementary Figure SF1**), which are not reflected by these F_v_/F_m_ values, and which were most probably a consequence of heat induced senescence (Li et al., 2021). In summary, these findings suggest that heat exposure to temperatures between 32 °C and 34 °C significantly increases the heat tolerance of the plasma membrane of *Arabidopsis thaliana,* which was not observed for higher temperatures. Heat induced effects on F_v_/F_m_ are robust across a wide range of acclimation temperatures which limits its applicability as heat stress indicator.

### Relative proportions of subcellular compartments and leaf tissues remain constant during heat acclimation

Heat can influence the architecture of plant cells and the different organelles within, varying depending on the duration and severity of the heat exposure. In chloroplasts, grana thylakoids were found to disorganise and unstack during heat exposure, especially at temperatures higher than 34 °C, and plastoglobuli numbers and sizes were found to increase with acclimation temperature (**Supplementary Figure S3**). This has also been reported before, together with swelling of chloroplasts and mitochondria due to heat exposure (Gounaris et al., 1984; Vani et al., 2001; Zhang et al., 2010; Grigorova et al., 2012; Zhang et al., 2014; Zou et al., 2017; Jampoh et al., 2023). The reconstruction of leaf cells based on SBF-SEM in the present study revealed constant relative proportions of chloroplasts (13 – 14 %), cytosol (3 - 4 %) and vacuole (82 - 83 %) before and after heat acclimation at 34 °C. In contrast, leaf disc volumes decreased by ∼ 30 % in heat acclimated plants while dry weight of the discs was not affected. This indicates a reduced (leaf) water content of heat acclimated plants which might be due to higher respiration rates under such conditions (Romero-Montepaone et al., 2021). Thus, while relative proportions of compartments remained constant, their absolute volumes decreased because of reduced total leaf tissue volume which suggests control of rather relative subcellular proportions than absolute compartment sizes. Similarly, also proportions of tissue types remained constant during heat acclimation (see Table II). Except for intercellular gas space, which decreased under heat, proportions of palisade mesophyll, spongy mesophyll, epidermal tissue and vascular bundles were similar before and after heat acclimation. In the present study, leaf cell structure was not analysed for plants which were less efficiently heat acclimated, e.g., after 7 days at 36 °C. This leaves room for speculation if the observed proportional reduction of (sub)cellular structures and leaf tissue composition is a prerequisite for or a consequence of heat tolerance. Yet, as membrane remodelling was proven earlier to be essential for efficient heat acclimation (Kunst et al., 1989; Murakami et al., 2000; Shiva et al., 2020), the proportional reduction of compartment size might be accompanied by, or even facilitate, remodelling processes, e.g., changes in saturation degrees of membrane lipids and classes, due to reduced physical distances between membrane systems. Particularly, for vesicle trafficking and membrane contact-based lipid transfer (Shomo et al., 2024), such a reduction of physical distance might be beneficial.

### Heat acclimation induces subcellular sugar allocation to stabilise sucrose metabolism during heat acclimation

A characteristic feature of eukaryotic cells is their compartmentation of metabolism which enables the separation and specification of pathways and their regulation (Hurry, 2017). Carbohydrates are direct products of chloroplast-located photosynthetic CO_2_ assimilation which, following the fixation and reduction reactions of the CBBC, are allocated to different subcellular compartments. This results in metabolite dynamics which are difficult to trace because transport processes across membranes systems and enzymatic interconversions are versatile (Sweetlove et al., 2017; Pommerrenig et al., 2018). The method of non-aqueous fractionation (NAF) enables the immediate and persistent quenching of enzymatic reactions which conserves the metabolic status at the sampling time point (Gerhardt and Heldt, 1984). In the present study, NAF revealed that, under heat, relative sucrose proportions are significantly decreased in the chloroplast and significantly increased in the vacuole. Together with the absolutely quantified metabolite amounts, this resulted in a significant increase of sucrose and glucose amounts in the vacuole. While such a vacuolar shift has also been reported before for cold acclimation (Knaupp et al., 2011; Fürtauer et al., 2016), the observed plastidial depletion of sucrose during heat acclimation contrasted findings made for low temperature (Nägele and Heyer, 2013). Together with members of the raffinose family oligosaccharides (RFOs), sucrose was shown to protect liposomes *in vitro* against damage by fusion which suggested also a protective function *in vivo*, e.g., under drought or freezing temperatures (Hincha et al., 2003; Knaupp et al., 2011). In contrast to cold or freezing, membrane fluidity is increased under heat which may not need the accumulation of sucrose or raffinose in the chloroplast to prevent fusion of thylakoid membranes. Instead, when subcellular sugar concentrations were calculated from total sugar amounts and estimated compartment volumes, cytosolic sucrose concentrations became highest under both 22 °C and 34 °C due to the comparatively low cytosolic volume. For glucose and fructose, a significant heat-induced cytosolic increase was observed which immediately raised the hypothesis of hexose accumulation due to affected invertase activity. Quantifying cytosolic neutral invertase activity revealed a stabilised activity between 22 °C and 34 °C, which became evident after normalisation to leaf tissue volumes and correction for thermodynamic effects applying the Arrhenius equation. In contrast, vacuolar invertase activity was significantly elevated by heat exposure. This led to the hypothesis that sucrose is transported along its concentration gradient from the cytosol to the vacuole where it is cleaved hydrolytically to release glucose and fructose. Such metabolite transport across the tonoplast might be facilitated by monosaccharide transporters as explained and outlined before (see e.g., (Pommerrenig et al., 2018)). The observation made in the present study that subcellular sugar concentrations, based on estimated compartment volumes, suggest a concentration gradient of sucrose from the cytosol into the vacuole of mesophyll cells might also indicate differential functions of proton-driven transport mechanisms which are currently thought to be necessary to drive metabolite exchange across the tonoplast along the proton gradient from vacuole into the cytosol.

Dynamics of both subcellular metabolite concentrations and enzyme activities result in dynamics of enzymatic reaction rates if metabolites act as substrates, products or regulatory effectors of enzymes. Simulation of reaction rates catalysed by neutral and acidic invertases, showed that, under heat, neutral invertase flux became very low which was caused by accumulation of hexoses in the cytosol which act as inhibitors (Sturm, 1999). While also acidic invertase flux was significantly affected by heat-induced accumulation of hexoses in the vacuole, a strong and significant increase of V_max_ counteracted this inhibition resulting in similar reaction rates at 22 °C and 34 °C (see Figure 8). Similarly, for plant cold acclimation, it was discussed earlier that temperature-induced shift of sucrose cleavage into the vacuolar compartment might stabilise metabolism and photosynthesis (Weiszmann et al., 2018). In summary, the presented findings suggest a conserved stabilizing role of vacuolar sucrose cleavage, and a specific role of sucrose accumulation and depletion in chloroplasts under low and elevated temperature, respectively.

## Author contributions

CS performed leakage and PAM assay, NAF analysis, microscopy, data analysis and wrote the paper. MH performed SBF-SEM microscopy. LS quantified enzyme activities. AK supervised and supported microscopy. TN conceived the study, analysed data, developed the R Shiny app and wrote the paper.

## Acknowledgements

We thank all members of Plant Evolutionary Cell Biology and Plant Development at LMU München for constructive discussions and advice. We also thank Kareem Farghaly, Keshav Goyal, Niklas Kroll, Marius Kröll and Suet Ying Wong for help with the annotation of the microscopy datasets. This work was supported by Deutsche Forschungsgemeinschaft (DFG, German Research Foundation) – INST86/1940-1 FUGG and TRR175/D03.

## Data availability statement

Data is provided in the supplements and, on request, by the corresponding authors. The code of a R Shiny app for kinetic simulations is provided on GitHub, https://github.com/cellbiomaths/Shiny_MM_simulation.

## Supplementary Legends

**Supplemental Table ST1. Settings for the “DL Training Segmentation 2D” module in Amira Pro for AI based segmentation.** The first training step was conducted according to the preliminary training instructions given in the Amira user’s guide without weights, all subsequent training steps used the previous results as weights. If settings were changed between trainings, a range of the provided values is given. The 22 °C dataset required 7 subsequent training steps, the 34 °C dataset required 15 subsequent training steps.

**Supplemental Table ST2. Data and calculation of subcellular volumes.**

**Supplemental Table ST3. Total and subcellular metabolite amounts, relative distributions and concentrations.**

**Supplemental Table ST4. Temperature correction of enzyme activities.**

**Supplemental Table ST5. Enzyme parameters for invertase simulations.**

**Figure SF1.**
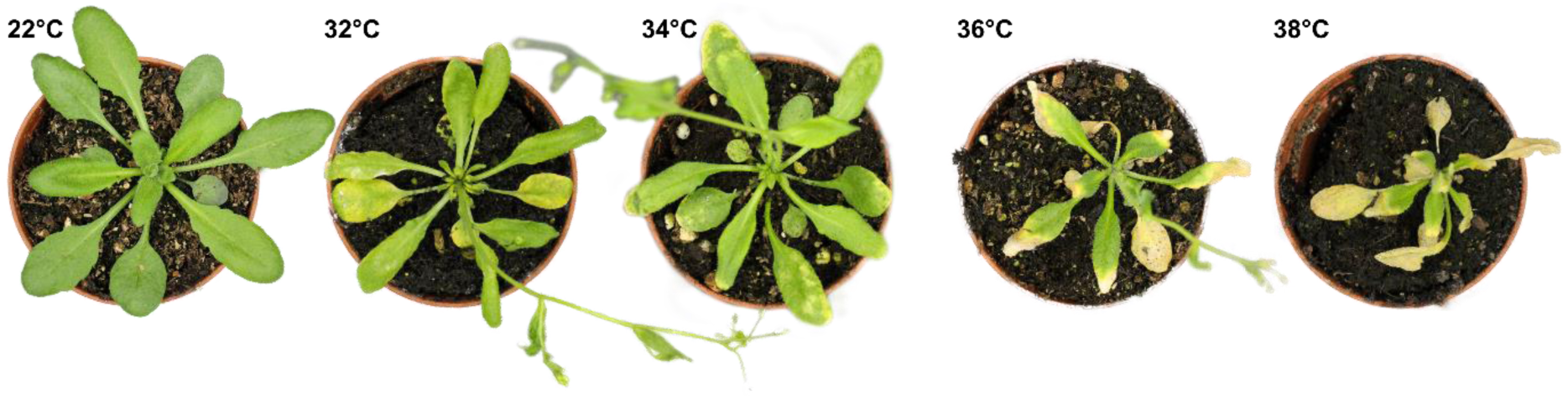
Plant phenotypes after heat treatment. From left to right: control (22 °C), 7 days 32 °C, 7 days 34 °C, 7 days 36 °C and 3 days 38 °C. Treatment with 32 °C to 36 °C results in yellowing of leaves in increasing severity and early induction of inflorescence. Treatment with 38 °C results in a low survival rate beyond 3 days of heat exposure. Surviving plants show heavy yellowing of leaves and no induction of inflorescence.

**Figure SF2.**
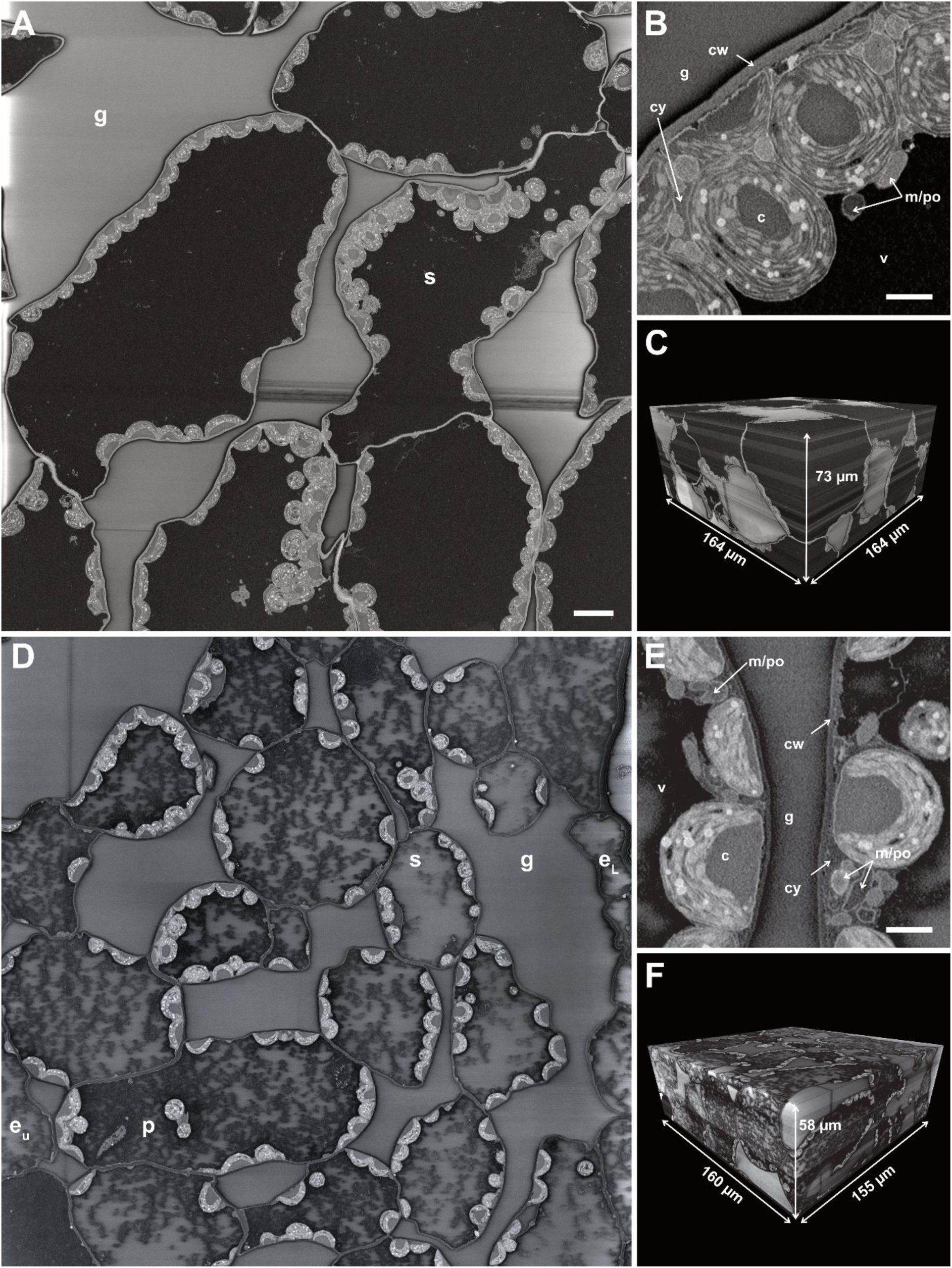
SBF-SEM of plant tissue. **A-C**: Control (22 °C), **D-F**: Heat treated (7 days 34 °C). **A+D**: Slice of sample block; e_U_ = upper epidermis, e_L_ = lower epidermis, g = gas space, p = palisade mesophyll cell, s = spongy mesophyll cell, scalebar = 10µm. **B+E**: High magnification of the same slice, c = chloroplast, cw = cell wall, cy = cytosol, g = gas space, m/po = mitochondria or peroxisome, v = vacuole, scalebar = 2µm. **C+F**: Complete sample block with dimensions.

**Figure SF3.**
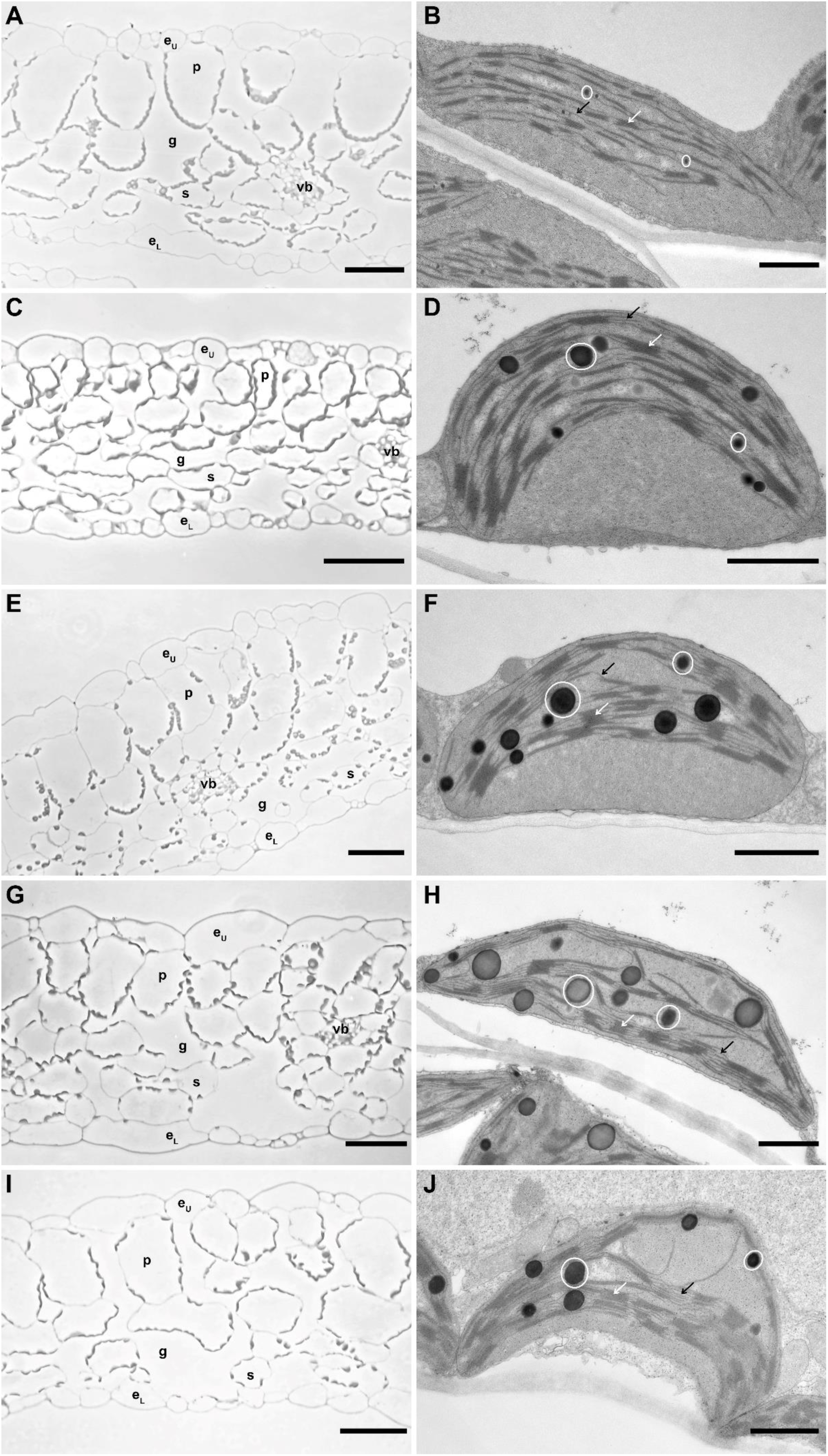
Light and electron micrographs of control and heat-treated leaves of *Arabidopsis thaliana*. **A, C, E, G, I:** Light microscopic images of semi thin sections of embedded leaf material; e_U_: upper epidermis, e_L_: lower epidermis, g: gas space, p: palisade mesophyll, s: spongy mesophyll, vb: vascular bundle, scalebar = 50 µm. **B, D, F, H, J:** Transmission electron micrographs of chloroplasts in ultrathin sections of embedded leaf material; white circles: plastoglobules, white arrows: grana thylakoids, black arrows: stroma thylakoids, scalebar = 1 µm. **A**: leaf section of control plant (22 °C). The leaf section shows typical architecture for *A. thaliana* leaves, the palisade mesophyll constitutes the upper layer of the mesophyll tissue, the spongy mesophyll the lower layer, vascular bundles are located in the middle, mostly in the spongy mesophyll; upper and lower epidermis both contain stomata and enclose the mesophyll tissue. **B:** Chloroplast of control plant (22 °C). The thylakoid membranes are arranged regularly and parallel to the vacuole facing side of the chloroplast, the plastoglobules are small and evenly distributed between the thylakoid membranes. **C:** Leaf section after 7 days at 32 °C; the tissue is packed more densely, and the cells are in average smaller than in the control leaf. **D:** Chloroplast after 7 days of 32 °C; the thylakoid membranes are still arranged similarly to the control, but the grana stacks are higher; the plastoglobules are larger than in the 22 °C chloroplast. **E:** Leaf section after 7 days of 34 °C; the leaf architecture is comparable to 32 °C, the tissue is more densely packed than in the 22 °C leaf. **F:** Chloroplast after 7 days of 34 °C; the thylakoid arrangement is comparable to 32 °C, however, the plastoglobules are larger than control and 32 °C. **G:** Leaf section after 7 days of 36 °C; the tissue is less dense than in the 32 °C leaves, but still denser, with smaller cells than in control plants, and some cells exhibit a less turgid shape compared to lower temperatures. **H:** Chloroplast after 7 days of 36 °C; the thylakoids are arranged less regularly and are disturbed by the large plastoglobules. **I:** Leaf section after 3 days of 38 °C; the density of the tissue is comparable to the control leaf, but some cells also exhibit a less turgid shape. **J:** Chloroplast after 3 days of 38 °C; the chloroplast has a crescent shape, compared to the more lentil-like shape of chloroplasts of lower temperatures; the thylakoids are arranged similar to the 36 °C thylakoids; the size of the plastoglobules is comparable to 34 °C.

